# Impact of Prior Cancer on Outcomes in Nasopharyngeal Carcinoma

**DOI:** 10.1101/542126

**Authors:** Huaqiang Zhou, Yaxiong Zhang, Jiaqing Liu, Wenfeng Fang, Yunpeng Yang, Shaodong Hong, Gang Chen, Shen Zhao, Jiayi Shen, Wei Xian, Zhonghan Zhang, Xi Chen, Hongyun Zhao, Yan Huang, Li Zhang

**Author notes:** Zhou, Zhang and Liu contributed equally to this work and should be regarded as co-first authors. **Correspondence information: Li Zhang** Mailing address: Department of Medical Oncology, Sun Yat–sen University Cancer Center, 651, Dongfeng Road East, Guangzhou, Guangdong, China, 510060. **Funding:** National Key R&D Program of China (2016YFC0905500, 2016YFC0905503), Chinese National Natural Science Foundation project (81572659).

## Abstract

**Background:** Prior cancer is a common exclusion criterion in nasopharyngeal carcinoma (NPC) trials. However, whether a prior cancer diagnosis affects trial outcomes is still unknown. We aimed to determine the impact of prior cancer on survival in NPC.

**Methods:** We identified patients diagnosed with NPC between 2004 and 2009 in the Surveillance, Epidemiology, and End Results (SEER) database. Variables were compared by chi-squared test and t-test as appropriate. Propensity score-adjusted Kaplan-Meier methods and Cox proportional hazard models were used to evaluate the impact of prior cancer on overall survival (OS).

**Results:** Among 3,131 eligible NPC patients, 349 (11.15%) patients had a history of prior cancer. The Kaplan-Meier curves did not show a statistically significantly different OS (*p*=0.19). Subgroup analyses stratified by timing of prior cancer and AJCC TNM stage of index cancer displayed the same tendency, prior cancer didn’t adversely affect OS compared with patients without prior cancer (*p*>0.05). Furthermore, in propensity score–adjusted COX models analysis, patients with prior cancer had the same/non-inferior OS (hazard ratio [HR] = 1.12, 95% confidence interval= 0.88 to 1.42).

**Conclusions:** Among patients with nasopharyngeal carcinoma, prior cancer does not convey an adverse effect on clinical outcomes, regardless of the timing of prior cancer and AJCC TNM stage of index cancer. Broader inclusion trial criteria could be adopted in nasopharyngeal carcinoma patients with a history of prior cancer. However, further studies are still needed to confirm.

## Introduction

Nasopharyngeal carcinoma (NPC) is a head and neck cancer common in south China and southeastern Asia(1). With the primary treatment of radiotherapy or chemoradiotherapy, the five-year overall survival (OS) of early stage nasopharyngeal carcinoma is greater than 90%(2). But recurrent or primary metastatic NPC still represents a critical unmet medical need in oncology research. Despite Intensity-modulated radiation therapy significantly improves the tumor local control rate, and distant metastasis is still poorly controlled, which remains the major reason of treatment failure. Many large clinical trials have been conducted to seeking the optimum comprehensive therapy to improve the survival, such as the standard first-line treatment option (gemcitabine plus cisplatin) and induction chemotherapy plus concurrent chemoradiotherapy for these patients(3,4).

Clinical trials are essential for better management of these patients. Fewer than five percent of adults with cancer in the United States participate in clinical trials(5). Clinical trial eligibility criteria present a major barrier to study enrollment, especially in oncology clinical trials, where patients with a prior cancer diagnosis are frequently excluded(6). For instance, over 80% of lung cancer trials sponsored by the Eastern Cooperative Oncology Group (ECOG) exclude patients with prior cancer(7). This practice is mainly based on the long-held assumption that prior cancer diagnosis and treatment could interfere with study outcomes. However, our previous pan-cancer study suggested that not all prior cancers actually interfere with study outcomes(8). The number of cancer survivors has a 4-fold increase in the United States over the last three decades(9). Due to the improved survival of cancer patients, the prevalence of multiple primary cancer also increases rapidly(10). Twenty-five percent of older adults and more than 10% of younger adults diagnosed with cancer have a history of prior cancer(11). Given the increased number of cancer survivors, the impact of this exclusion criteria will likely increase.

Until now, no study has specifically evaluating the impact of prior cancer on NPC outcomes, and little is known about the characteristics of NPC patients with prior cancer. To address these assumptions, we therefore determined the characteristics and prognostic impact of prior cancer among patients with nasopharyngeal carcinoma using the Surveillance, Epidemiology, and End Results (SEER) database.

## Methods

### Data source and population

We extracted data from the SEER database by using the SEER*Stat software version 8.3.5, which covers approximately 30% of the population in the United States (https://seer.cancer.gov/, accession number: 13693-Nov2015)(12,13).

The study population included patients diagnosed with NPC from January 2004 to December 2009. Patients who meet any of the following criteria were excluded from the study: (1) age at diagnosis younger than 18 years; (2) patients with only autopsy or death certificate records; and (3) patients with incomplete survival data and follow-up information.

We extracted demographic and clinicopathological data from SEER database, including sex, age, race, marital status, pathology grade, TNM stage, surgery, and radiotherapy. We classified race as white, black and others. Patients were divided into married or unmarried. The TNM stage was based on the AJCC (6th edition) staging system. Considering that the survival data were available in the measurement unit of months, a survival time of 0 month was recorded as 0.5 months to include patients who died within one month of diagnosis.

### Measures

A history of prior cancer was determined from SEER sequence numbers, as described in our previous study(8). In brief, sequence numbers represent the order of all primary reportable neoplasms diagnosed in a lifetime. The timing of the prior cancer was calculated by using the SEER diagnosis dates of the index cancer and the most recent of any prior cancers. Cases with full timing records were used for further study. The primary outcome of this study was overall survival. We set December 31, 2014 as the follow-up cutoff date to ensure that all included cases were followed up for at least 5 years.

### Statistical analysis

We categorized patients into two groups based on prior cancer history. Differences in patients’ characteristics were assessed by Pearson chi-squared analysis for categorical variables and t-test for continuous variables as appropriate. In this study, we employed a propensity score matching (PSM) method to minimize the effect of confounding from differences in baseline characteristics(14). Propensity scores were calculated based on race, sex, age, marital status, TNM stage, pathologic grade, and treatment. A one-to-one PSM with a caliper of 0.2 was performed. The characteristics were balanced after PSM. These PSM pairs were used in subsequent analyses.

OS was calculated with the Kaplan-Meier method, and differences were compared using log-rank tests. Finally, we built a multivariate Cox proportional hazards model to identify whether prior cancer impacted the prognosis independently. The common demographic and clinicopathological data, including race, sex, age, marital status, TNM stage, pathologic grade and treatment, were entered as covariates. Statistical significance was set as a two-sided p-value less than 0.05. Analyses were performed using R Statistical software (version 3.4.2, Institute for Statistics and Mathematics, Vienna, Austria; www.r-project.org).

## Results

In total, we identified 3,131 eligible NPC patients diagnosed between 2004 and 2009. Among these cases, 11.15% (n=349) had a history of prior cancer. Compared with cases without previous malignancies, patients with prior cancer were older (66.25 vs 54.59 years, *p*<0.001), female (37.0% vs 29.5%, *p*=0.005), white (70.5% vs 48.5%, *p*<0.001), and unmarried (47.3% vs 40.7%). The percentage of surgery was larger among patients with a prior cancer (15.8% vs 11.9%, *p*=0.047), and patients with prior cancer received less radiotherapy (67.9% vs 80.1%, *p*<0.001). Additional baseline characteristics are displayed in Table 1. Characteristics were balanced between groups after adjustment for propensity score (Table 1, *p* > 0.05). Among 349 NPC patients with a history of cancer, prostate (20.60%), gastrointestinal (13.73%), other genitourinary and gynecologic (12.88%), and breast cancers (9.44%) were the most common types of prior cancer. Localized and regional stages were accounting for 77.46% of cases. Over 60.52% of prior cancer were diagnosed within 5 years of the index NPC. The median time between the most recent prior cancer diagnosis and the index NPC was 3.5 years.

**Table 1.**
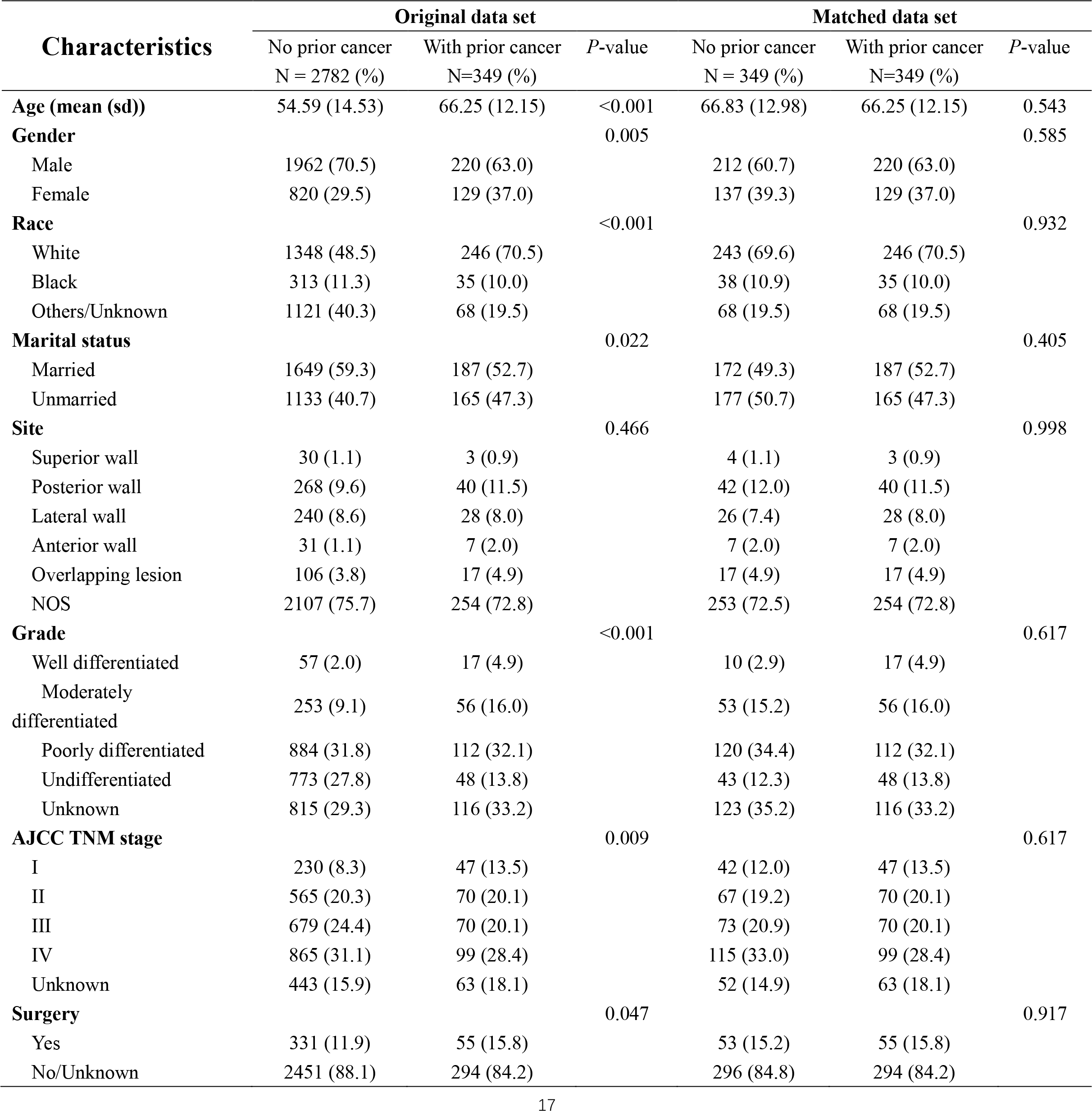
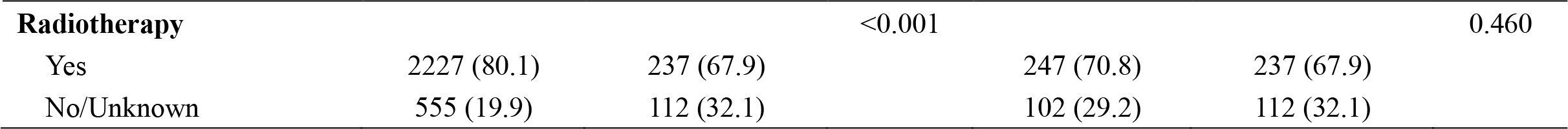
Baseline characteristics of patients with nasopharyngeal carcinoma in the original/matched data sets (N=3131).

In unadjusted Kaplan–Meier analysis, NPC patients with prior cancer demonstrated similar OS compared to patients without prior cancer (log rank tests *p*=0.19) (Figure 1). The overall five-year survival rate for patients with or without prior cancer were 35.2% (95% confidence interval (CI), 30.5–40.6) and 39.8% (95% CI, 35.0–45.4), respectively. Figure 2 depicts Kaplan–Meier survival curves stratified by the timing of prior cancer and index cancer TNM stage. Subgroup analyses stratified by timing of prior cancer and AJCC TNM stage of index cancer displayed the same tendency, prior cancer didn’t adversely affect OS compared with patients without prior cancer (*p*>0.05).

**Figure 1.**
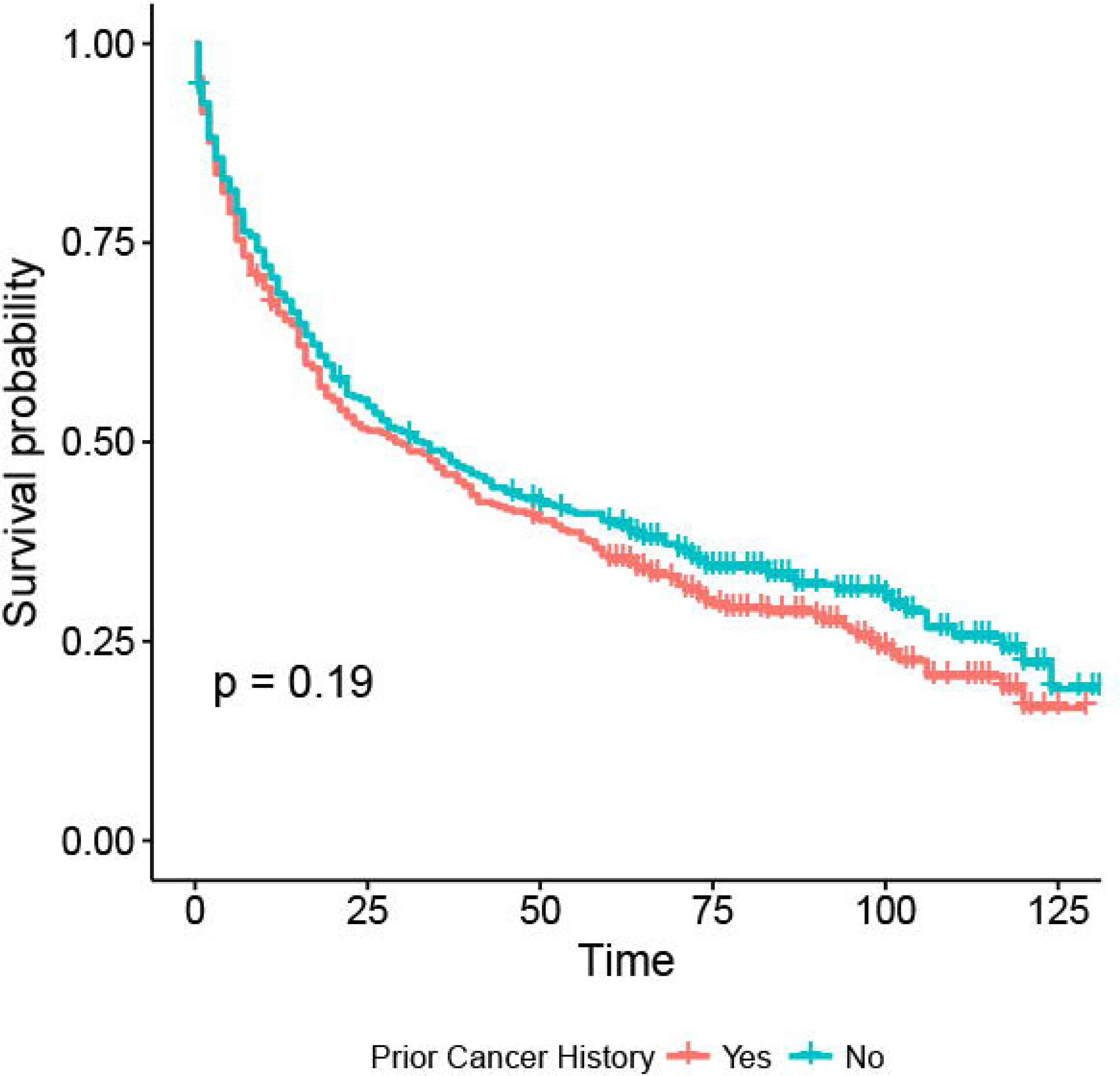
The Kaplan-Meier survival curves of prior cancer impact on the overall survival in nasopharyngeal carcinoma. The overall survival of nasopharyngeal carcinoma was similar compared with that of patients without a prior cancer (*p*>0.05).

**Figure 2.**
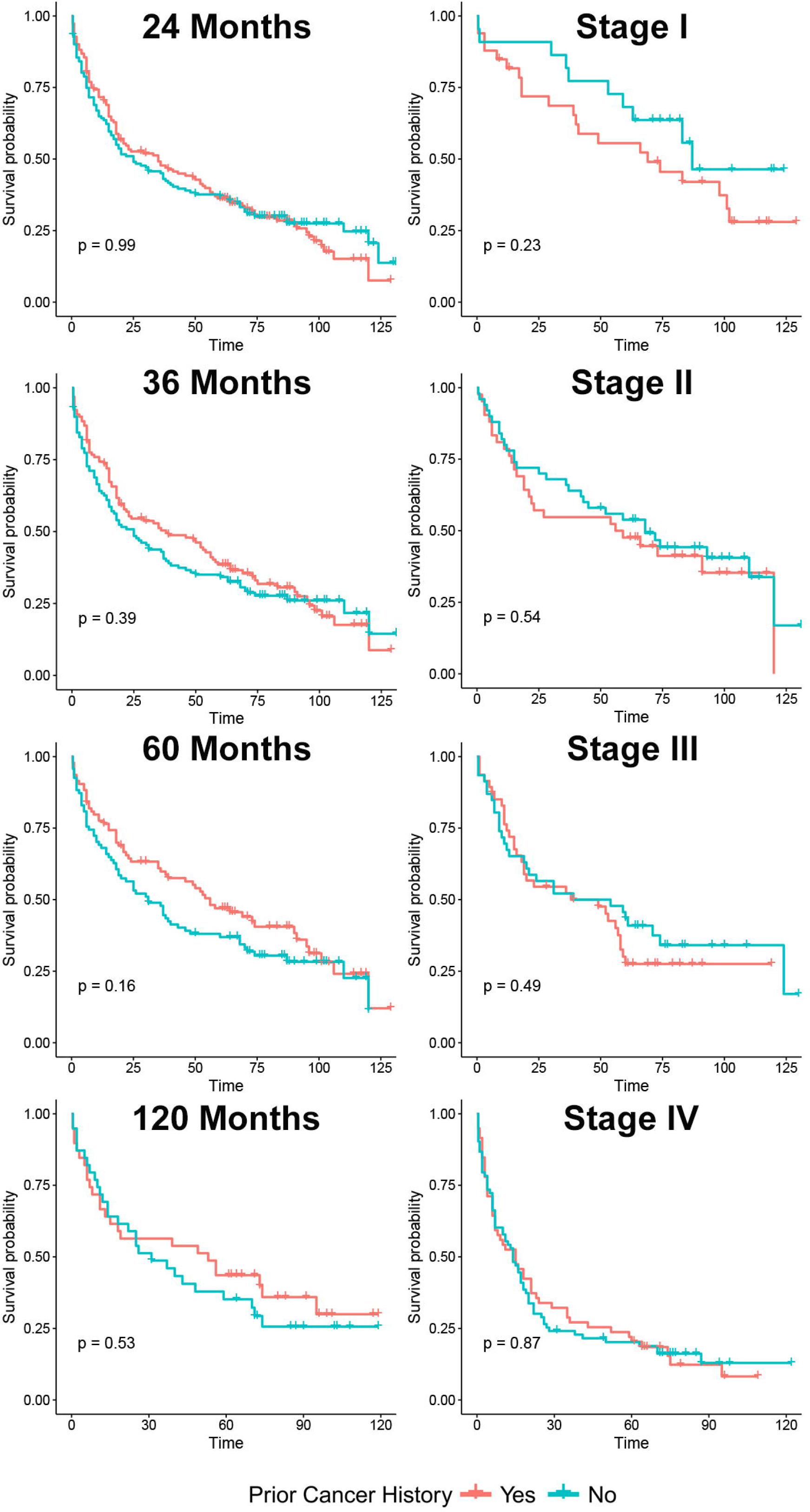
Subgroup analysis of prior cancer impact on the overall survival stratified by the timing of prior cancer and AJCC stage in nasopharyngeal carcinoma. The nasopharyngeal carcinoma patients with prior cancer showed a similar survival when compared with patients with no prior cancer, regardless of the timing of prior cancer and stage.

In propensity-score–adjusted Cox models, patients with prior cancer had the same/non-inferior OS (hazard ratio [HR] = 1.12, 95% confidence interval= 0.88 to 1.42), compared to patients without a prior cancer (Table 2).

**Table 2.**
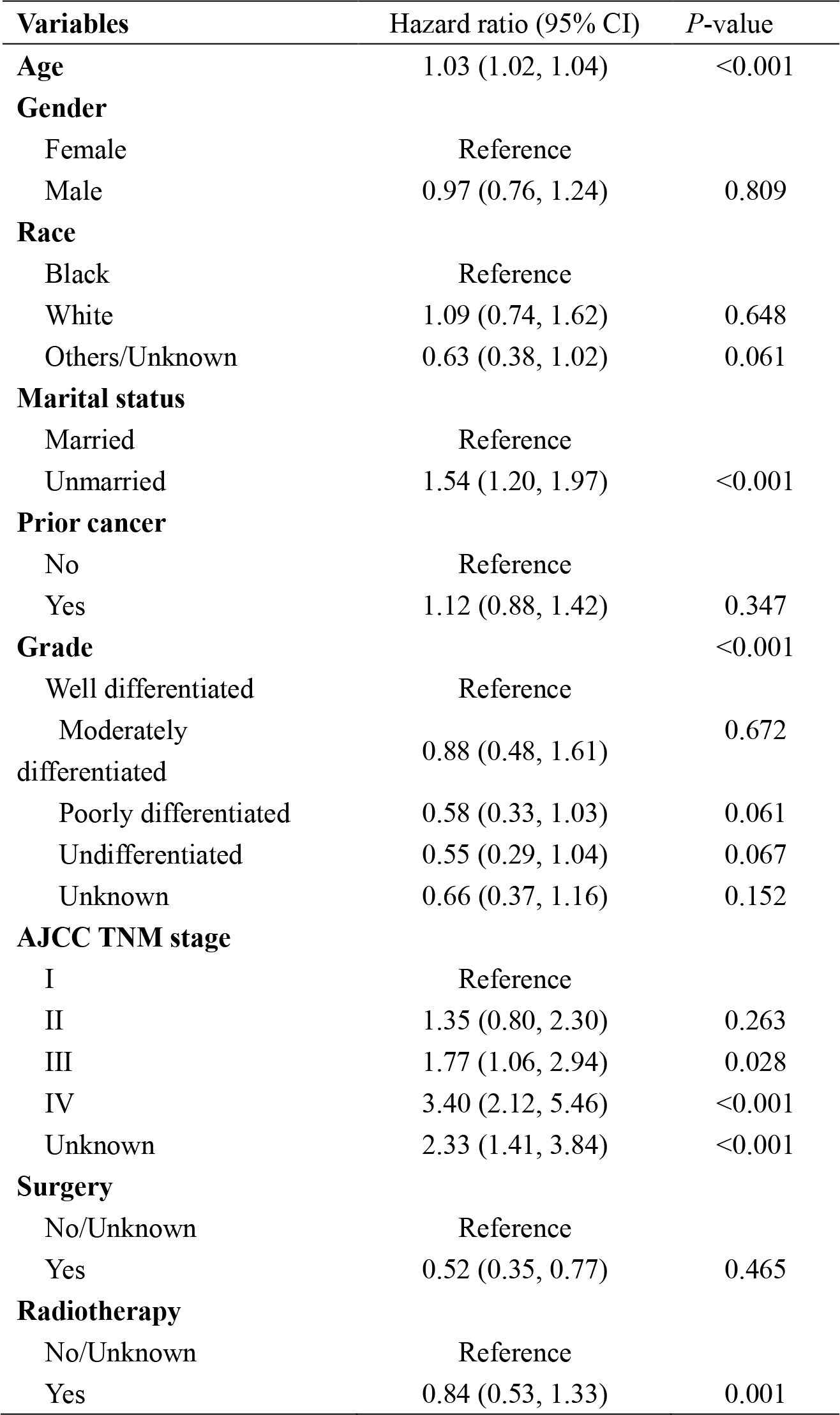
Cox regression analysis of prior cancer history impact on patients with nasopharyngeal carcinoma.

## Discussion

Stringent eligibility criteria for oncology clinical trials can minimize the risks to the participants, but they can affect the accrual and external validity of clinical trial significantly(15). In practice, patients with a prior cancer history are usually excluded in cancer clinical trials due to the potential interference of study outcomes. But there is no authoritative data to this assumption now. Given the sizeable number of cancer survivors, the impact of this criteria will increase, and it is critical to understand the impact of prior cancer. Up to now, whether NPC patients with prior cancer faced a worse prognosis remains unknown. Our study fills this knowledge gap exactly. We observed that NPC patients with prior cancer did not result in inferior survival outcomes when compared with those without prior cancer. To our knowledge, this is the first study to evaluate the characteristics and prognostic impact of prior cancer among NPC patients. Thus, we need to rethink the long-held practices that patients with prior cancer are excluded from clinical trials.

Our previous study has mentioned the different impact of prior cancer according to the specific cancer(8). We creatively divided these cancers into two categories, “prior cancer inferior” (PCI), in which patients had lower survival rates than those without prior cancer; and “prior cancer similar” (PCS), in which survival rates were similar. From this point of view, NPC is one kind of PCS cancers. Several studies also addressed the same questions for other cancer types. Although prior cancer might impact the overall survival in patients with prostate cancer(16), prior cancer does not contribute to poor survival outcome in many other cancer types, such as lung, glioblastoma, esophageal, gastrointestinal tract, pancreatic cancer(17–23). Notably, the impact of prior cancer on early-stage, locally advanced and advanced lung cancer are consistent, without adverse effect on clinical outcomes(17,18,20). Our results also confirmed similar phenomena in different stage of NPC for the first time. It suggests that our findings are applicable to clinical trials for different stages of NPC.

The timing of prior cancer is also needed to be fully considered when determining the impact of prior cancer exclusion criteria on clinical trials(24). Generally, a 5-year exclusion window is commonly employed in most trials(7), and over 60% of prior cancers occur within this window in our study. The median interval between a prior cancer and the index NPC was 3.5 years. This information indicated that active surveillance and screening for NPC is necessary in cancer survivors. Subgroup analysis stratified by timing of prior cancer displayed the same tendency, prior cancer did not adversely affect OS. In other words, the impact of prior cancer is independent of timing. From this perspective, NPC patients with prior cancer can be considered for enrollment in trials regardless of timing, and improve accrual without affecting outcomes.

However, there are several limitations in interpreting our study. Firstly, there is a paucity of detailed characteristics about prior cancers diagnosed outside of the registry state, which are recorded in sequence number only. So, our study only focused on the timing of the prior cancer. Additionally, the efficacy and tolerability of therapy on prior cancer, which may disrupt the management for the index cancer, cannot be considered due to the data restriction. Secondly, we cannot obtain detailed data on treatments and comorbidities from the SEER database. Therefore, neither did comorbidities be matched in our PSM analyses, nor did they are included in the regression models. Thirdly, PSM analysis only accounts for observable covariates, and hidden bias resulted from unobserved confounders remain after matching. Finally, the data obtained from SEER cover approximately 34.6% of the total U.S. population, thus making it necessary to confirm the generality of our findings.

## Conclusions

Among patients with nasopharyngeal carcinoma, prior cancer does not convey an adverse effect on clinical outcomes, regardless of timing of prior cancer and stage of index cancer. Broader inclusion trial criteria could be adopted in nasopharyngeal carcinoma patients with a history of prior cancer. However, further studies are warranted to confirm the appropriateness of this exclusion criterion in nasopharyngeal carcinoma trials.

## Abbreviations

NPC: Nasopharyngeal carcinoma
OS: Overall survival
ECOG: Eastern Cooperative Oncology Group
SEER: Surveillance, Epidemiology, and End Results
PSM: Propensity score matching
CI: Confidence interval
HR: Hazard ratio
PCI: Prior cancer inferior
PCS: Prior cancer similar

## Declarations

### Ethics approval and consent to participate

Institutional review board approval was waived for this study because SEER database is a public anonymized database. The author Zhou. has gotten the access to the SEER database (accession number: 13693-Nov2015).

### Consent for publication

All authors have confirmed the manuscript and approved the publication of the manuscript. The corresponding author has completed the “Consent for publication”.

### Availability of data and materials

The datasets generated and/or analysed during the current study are available in the SEER database, (https://seer.cancer.gov/).

### Competing interests

All of the authors have no conflicts of interested to declare.

### Funding

National Key R&D Program of China (2016YFC0905500, 2016YFC0905503), Chinese National Natural Science Foundation project (81572659).

### Authors’ contributions

L. Zhang, H.Q. Zhou, Y.X. Zhang and J.Q. Liu were responsible for the conception and design of the study, interpretation of data, drafting, and writing of the article; W.F. Fang, Y.P. Yang, S.D Hong, G. Chen, and S. Zhao were responsible for acquisition analysis and interpretation of data, drafting the text and also participated in the drafting of the article; J.Y. Shen, W. Xian, Z.Z Zhang and X. Chen were responsible for interpretation of data and drawing figures. H.Y. Zhao and Y. Huang were responsible for revision of the intellectual content. All authors participated in final approval of the article and agreed to be accountable for all aspects of the work.

## Acknowledgements

The authors acknowledge the efforts of the Surveillance, Epidemiology, and End Results (SEER) Program tumor registries in providing high quality open resources for researchers.

